# Thermal tolerance of mosquito eggs is associated with urban adaptation and human interactions

**DOI:** 10.1101/2024.03.22.586322

**Authors:** Souvik Chakraborty, Emily Zigmond, Sher Shah, Massamba Sylla, Jewelna Akorli, Sampson Otoo, Noah H. Rose, Carolyn S. McBride, Peter A. Armbruster, Joshua B. Benoit

## Abstract

Climate change is expected to profoundly affect mosquito distributions and their ability to serve as vectors for disease, specifically with the anticipated increase in heat waves. The rising temperature and frequent heat waves can accelerate mosquito life cycles, facilitating higher disease transmission. Conversely, higher temperatures could increase mosquito mortality as a negative consequence. Warmer temperatures are associated with increased human density, suggesting a need for anthropophilic mosquitoes to adapt to be more hardy to heat stress. Mosquito eggs provide an opportunity to study the biological impact of climate warming as this stage is stationary and must tolerate temperatures at the site of female oviposition. As such, egg thermotolerance is critical for survival in a specific habitat. In nature, *Aedes* mosquitoes exhibit different behavioral phenotypes, where specific populations prefer depositing eggs in tree holes and prefer feeding non-human vertebrates. In contrast, others, particularly human-biting specialists, favor laying eggs in artificial containers near human dwellings. This study examined the thermotolerance of eggs, along with adult stages, for *Aedes aegypti* and *Ae. albopictus* lineages associated with known ancestry and shifts in their relationship with humans. Mosquitoes collected from areas with higher human population density, displaying increased human preference, and having a human-associated ancestry profile have increased egg viability following high-temperature stress. Unlike eggs, thermal tolerance among adults showed no significant correlation based on the area of collection or human-associated ancestry. This study highlights that the egg stage is likely critical to mosquito survival when associated with humans and needs to be accounted when predicting future mosquito distribution.

## Introduction

Mosquitoes pose a significant public health threat due to their ability to transmit a wide range of pathogens, including Zika, chikungunya, and West Nile viruses, leading to substantial morbidity and mortality worldwide (Merle et al., 2018; Shragai et al., 2017). Among the various mosquito-borne diseases, dengue fever, a disease caused by a viral infection transmitted by *Aedes* mosquitoes, has more than quadrupled over the last decade (Bhatt et al., 2013; Brady et al., 2012). Another mosquito-borne virus, Zika virus, has emerged rapidly since 2007, giving rise to epidemics in regions such as Micronesia, the South Pacific, and most notably, the Americas (Weaver et al., 2016). While the primary vector for dengue and Zika is *Ae. aegypti*, *Ae. albopictus* can also transmit these viruses (Lai et al., 2020; Lambrechts et al., 2010; McKenzie et al., 2019; Rezza, 2012). The yellow fever mosquito, *Ae. aegypti* is a global invasive warm-weather species and thrives in urban and rural habitats across the Asian, African, and American tropics (Christophers & Others, 1960). There are records of *Ae. aegypti* are found across wild habitats of Africa, and these mosquitoes prefer to feed on forest animals (McBride et al., 2014). The globally distributed *Ae. aegypti* mosquitoes, mainly the ‘domestic’ form, evolved to bite humans and are the primary vector of dengue, chikungunya, and yellow fever. However, in the African landscape, these two types - the ‘domestic’ and the ‘wild’ still coexist in nature (McBride et al., 2014). These two subspecies of *Ae. aegypti* are *Ae. aegypti aegypti* (*‘Aaa’*), the domestic form with a strong affinity for human hosts, and *Ae. aegypti formosus* (‘*Aaf*’), the more versatile, generalist form with a broader range of hosts, including non-human animals (McBride et al., 2014; Rose et al., 2020). Another *Aedes sp.*, *Ae. albopictus*, the Asian tiger mosquito, is a highly adaptable species first described in Southeast Asia. Over the last two decades, *Ae. albopictus* has spread to Africa, the Middle East, North and South America, and the Caribbean (Bonizzoni et al., 2013).

Over the last 20-40 years, behavioral shifts, including host preferences and oviposition behavior, have been documented for *Ae. aegypti* and *Ae. albopictus*. At least partially, these behavioral shifts can be attributed to anthropophagy and breeding in human-associated environments (Ponlawat & Harrington, 2005; Richards et al., 2006; Rose et al., 2023). Multifaceted influences, such as changes in land use patterns (Swei et al., 2020), alteration of ecosystems (Vora, 2008), deforestation (Ortiz et al., 2021), urbanization (Esch et al., 2014; Melchiorri et al., 2018), and climate change (McBean & Ajibade, 2009; Reinmann et al., 2016) have led to the expansion of human settlements into previously uninhabited areas with favorable environmental conditions for mosquito breeding (Chala & Hamde, 2021). Densely populated human areas and unplanned urbanization (Kolimenakis et al., 2021; Moore et al., 2003), in conjunction with the heat rise due to anthropogenic effects (Akhtar et al., 2016; Mathieu & Karmali, 2016), likely promote the evolution of anthropophagy. Additionally, dry conditions force mosquitoes to seek out water for breeding and hydration (Benoit et al., 2023; Holmes & Benoit, 2019), such as during dry seasons where water near human residences may be the only sites to deposit eggs (Chumsri et al., 2018; Dhiman, 2016; Rose et al., 2023). The ovipositional and environmental conditions associated with high human density underscore the critical need for understanding mosquito biology in relation to rural vs. urban transitions and varying human preference.

To understand mosquito biology in different environmental contexts characterized by variations in human density, a useful starting point is to explore the significance of the egg stage. The mosquito egg stage is crucial for *Aedes* as embryos within the egg chorion can remain viable for extended periods of unfavorable conditions until water is available (Sota & Mogi, 1992). *Aedes* mosquitoes lay their eggs on wet surfaces near water sources and will subsequently emerge when the eggs become submerged in water (Clements, 1992; Day, 2016). Since mosquito eggs are immobile, they are particularly vulnerable to shifting environmental conditions. Among mosquito species, *Aedes* eggs show extraordinary resilience to abiotic stressors, especially dehydration, due to unique aspects of their embryonic development and properties of the egg cuticle (Farnesi et al., 2017; Faull et al., 2016; Hinton, 1968; Juliano et al., 2002; Rezende et al., 2008, 2016; Vargas et al., 2014). Another critical abiotic component that can impact the viability of *Aedes* eggs is fluctuations in temperature (Faull & Williams, 2015; C. L. Judson, 1960). Heat stress can lead to negative consequences, including increased egg and larvae mortality, reduced insect larval growth, and even impacts on subsequent adult stages (Ajayi et al., 2023; Bowler & Terblanche, 2008; Rocha et al., 2017). For example, temperatures exceeding 30°C negatively impacted the viability of *Anopheles gambiae* mosquito egg development (Goltsev et al., 2009; Impoinvil et al., 2007). Ovipositional choices for *Aedes* encompass natural locations like tree holes or artificial containers, mostly with darker hues like discarded car tires, and exposure to sunlight could drastically impact the thermal exposure of the eggs. The darker-colored artificial containers are subject to warming up a few degrees more compared to the shaded tree holes covered by a leafy canopy. In urban environments, where pollutants and artificial containers are abundant, *Aedes* mosquitoes frequently lay on surfaces that absorb heat readily and can retain higher temperatures for more extended periods. This starkly contrasts eggs laid in natural habitats and benefitted from a buffered environment. Furthermore, increased urbanization and higher human populations create urban heat islands (UHI) (Hsu et al., 2021; Tuholske et al., 2021). However, little is known about whether differences in the thermal tolerance of mosquito eggs are related to variation in human association. Here, we test the hypothesis that mosquito eggs from densely populated areas and increased human preference have a higher thermal tolerance capacity.

Previous mosquito egg viability and embryo development studies have primarily focused on the process of cuticular formation, factors such as oxygen concentrations that stimulate hatching, and interspecific variation in desiccation resistance (Faull & Williams, 2015; Judson & Hokama, 1965; Judson, 1960; Sota & Mogi, 1992). A comprehensive examination of habitat-specific egg thermal tolerance has yet to be conducted. The comparison of egg thermal tolerance in mosquitoes from human-associated compared to sparsely populated environments could serve as a catalyst for comprehending the adaptive egg thermal plasticity of *Aedes* eggs. We examined the impact of temperature stress on *Ae. aegypti* and *Ae. albopictus* eggs and adults. For *Ae. aegypti*, we examined the effects of host preference, genomic ancestry, and several other ecological indices as potential factors contributing to egg thermal tolerance based on recently collected field populations described in (Rose et al., 2020). The major highlight is a higher thermal tolerance of *Ae. aegypti* eggs, but not the adults, as the association with humans increases.

## Materials and Methods

### Mosquito collection and maintenance

*Aedes aegypti* mosquito colonies were established from eggs collected across Western Africa using seed germination papers in ovitraps (Rose et al., 2020). For our study, colonies from five locations in Senegal and two in Ghana were selected (**Supplementary table 1**). *Ae. albopictus* mosquito colonies were also established from eggs collected from the USA and Japan (**Supplementary table 1**) (Mushegian et al., 2021). The mosquito larvae were reared in plastic pans with a surface area of approximately 1500 cm^2^, maintaining a uniform density of approximately 0.165g/cm^3^ for the larvae. The rearing medium consisted of deionized (DI) water, finely ground fish food (Tetramin, Goldfish Flakes), and yeast extract (Difco, BD 210929). Mosquitoes were maintained in a climate-controlled facility (27±2 °C; 65-70% RH; 16-8-hour (hr) light-dark cycle). The adults were kept in 12″ x 12″ x 12″ mesh cages with access to water and 10% sucrose solution ad libitum. Females (12-14 days old) were blood-fed with a human host (University of Cincinnati IRB 2021-0971), and eggs were collected in brown hardwound towels for thermal stress studies. Experiments were conducted on eight (8) populations of *Ae. aegypti* and six (6) populations of *Ae. albopictus*, including a laboratory-maintained population from Florida (Benzon Research) for *Ae. aegypti* and a laboratory-maintained population from New Jersey (BEI Resource, NR-48979) for *Ae. albopictus* (**Supplementary table 1**). Additionally, a hybrid population of *Ae. albopictus* labeled “Combined” was formed with equal numbers of males and females from populations (SAG, JAC, BRU, and F-36) (**Supplementary table 1**). The thermal tolerance of eggs and adults was assessed for each population (described below). Due to the more extended periods that *Ae. albopictus* has been in the lab, and because of a lack of ecological conditions, human population levels and metrics related to human preference known at collection sites, models were not used to investigate this species in relation to human association.

### Egg thermal tolerance

Egg batches collected from each cage were held under colony conditions for two weeks to allow embryonation (Maïga et al., 2017). Eggs (n=30-50 per replicate on oviposition paper from multiple females with 10-12 replicates per treatment) were stored in 0.28 oz. size plastic vials [Thornton Plastics: 2.5UDbl]. Heat stress [25±1, 29±1, 33±1, 37±1, 41±1, and 45±1 °C] was generated using a digital dry bath (Thermo Scientific™). Plastic vials were placed in dry heat blocks and covered with insulated styrofoam to ensure a constant temperature. The initial temperature in the dry heat bath was 29°C. The experimental temperature increased to the treatment level for up to 30 minutes at approximately 0.25 °C per minute. A digital thermometer (OmegaTM) was used to confirm the temperature during each assay. Eggs were tested at 2 or 6 hr exposure for all experimental temperatures. After the temperature returned to 29°C, the egg vials were filled with DI water and fish food. Two days following the experiment, 80% ethanol was used to preserve larvae and eggshells. Vials were stored at -20°C until viability counts. During the egg counting process, we counted four stages/indices: hatched eggs, unhatched eggs, deformed eggs, and larvae. Hatched eggs were identified by the presence of a pore, slit, or broken cap, indicating the emergence of the larvae. The egg hatching rate was determined as the ratio of hatched eggs to the total number of eggs, which quantitatively measures the proportion of eggs that successfully hatched. Hatching rates, when eggs were exposed to control conditions, was over 92% for all populations, indicating that conditions to allow larval emergence were not a major factor in these studies, as previously observed (Metz et al., 2023).

### Adult thermal tolerance

Adults (males and females) were exposed to a range of temperatures - 33±1, 37±1, and 41±1 °C for 2 hr and 6 hr. Before the experiments, the adults were held at a climate-controlled facility (described previously) with access to water and 10% sucrose solution ad libitum. Eight 7–10-day old adult mosquitoes, separated by sex, were held in 50 ml perforated plastic vials to allow for airflow. The experiments were conducted with five replicate vials for each sex and temperature. Vials were partially submerged in a water bath (ThermoFisher Scientific) and covered with polystyrene foam to maintain temperature. The water bath started at 29 °C and gradually increased to the testing temperature at a rate of 0.25°C per minute. The temperature in the vials was verified with a digital thermometer (OmegaTM). All the trials were consistently conducted at approximately the same circadian time every day (9.00-10.30 am local time), and post-experiment, mosquitoes were moved to rearing conditions (described previously) for recovery. Survival was examined immediately (early: 2 hr) and later (24 hr) after the thermal stress by counting the mobile mosquitoes. Proportion survival was calculated as the ratio of mobile mosquitoes to the total mosquito number.

### Statistics and ecological modeling in relation to thermal stress

The deviation from average adult survival for each of the populations was plotted in heatmaps using the ‘pheatmap’ package in R. Similarly, deviation from average egg hatching for each population was plotted using dot plot. Differences in survival rates and egg viability were assessed using a linear regression model, ‘lm function’ in R (Chambers & Hastie, 1992), followed by a post-hoc comparisons test using the "emmeans" package (default -multiple comparison Tukey methods) in R (version 4.2.3), where *p* < 0.05 were considered statistically significant (Searle et al., 1980).

We examined the influence of ecological, behavioral, and ancestry indices (Rose et al., 2020) on the survival of adults and eggs following temperature stress. From these analyses, we contrasted the egg thermotolerance for mosquito populations from urban (areas with high human density) and rural (human areas with lower population) habitats and established a comprehensive evaluation based on ancestry components, mosquito preference, and human population density. In addition, we combined the ecological (human population density), behavioral (host preference), and genomic (ancestry) factors into a combined index (human specialization syndrome) framework to explain egg thermotolerance in the linear models described below. A three-step analysis (a correlation test, a binomial logistic regression, and a linear model) was used. In the first step, we employed a correlation analysis, utilizing a ‘cor’ test (default: ‘Pearson’) in R (Becker et al., 1988), to determine the relationships and dependencies among the behavioral and ecological indices. This comprehensive exploration of the variables aimed to evaluate the distinct specialization and adaptation strategies exhibited by two different subspecies of *Ae. aegypti,* i.e., the human specialist domestic form *‘Aaa’*, and the versatile, generalist form ‘*Aaf*’ (McBride et al., 2014; Rose et al., 2020). This exciting dichotomy plays a pivotal role in shaping the behavioral adaptations of *Ae. aegypti*, their interactions with humans, and vector-disease dynamics (Aubry et al., 2020; Bennett et al., 2021; McBride et al., 2014). In the second step, following the correlation test, we aimed to determine the variables that significantly influence egg hatching, and then the relationships between egg hatching and the significant variables were explored using a linear model framework. A binomial logistic regression modeling approach was used to estimate the significant variables influencing the egg-hatching success of the mosquito populations based on multiple factors, including temperature, stress periods/levels, preference for the host, ancestral origin associated with human specialization, precipitation, warm quarter precipitation, and human density of the locality of collection. These individual analyses were followed with a combined model that assessed all the factors listed before to assess for potential interactive effects. Lastly, in the third step, we estimated the impact of 2- and 6-hr-long temperature stresses on the hatching success with several other ecological and genomic parameters using a linear model framework that resulted in a coefficient for each mosquito collection location. In this set of analyses, each population was represented by a single regression factor (generated using linear modeling), and the data was scaled using the ‘scales’ package (Wickham & Seidel, 2016) in R to correlate with mosquito hatching success.

## Results

### Temperature tolerance of adult mosquitoes

In general, adult females showed increased thermal tolerance compared to adult males (*Ae. aegypti*: F = 51.76, *p* = 1.27 × 10^-16^; *Ae. albopictus*: F = 34.3, *p* = 7.2 × 10^-9^) and survival decreased as temperature increased (*Ae. aegypti*: F = 1679, *p* < 2.2 × 10^-16^; *Ae. albopictus*: F = 909.7, *p* < 2.2 × 10^-16^). For *Ae. aegypti,* we observed only a few significant differences in survival among the populations for males or females when compared to the same sex at each temperature treatment (33±1, 37±1, and 41±1 °C) (**Fig. 1; Supplementary tables S2a and S2b**). Survival was assessed at both 2 hr and 24 hr after the temperature treatment (**Fig 1; Supplementary figures 1a, 1b, 1c, and 1d**), and as expected, mortality increased with the longer elapsed treatment time (F = 44.69, *p* = 3.91 × 10^-11^). Similar responses were observed in *Ae. albopictus*, with few significant differences detected among each sex within the temperature-treated (33±1, 37±1, and 41±1 °C) for different populations (**Fig. 2; Supplementary figures 1e, 1f, 1g, and 1h; Supplementary tables S2c and S2d**). We observed lower survival of adult *Ae. albopictus* than *Ae. aegypti* (F = 23.786, *p* = 1.98 × 10^-6^; **Supplementary figures 2a and 2b**). Overall, females are more tolerant to heat stress than male mosquitoes for both species and survival decreased with increasing temperature (**Supplementary figure 3, Supplementary table S3a**), with only a few significant differences in adult survival between populations that do not have specific trends (**Supplementary tables S2a and S2c)**. Of interest, there was a trend for increased heat tolerance for lines from urban areas (NGO, THI, and KUM), but this was not significant.

**Figure 1:**
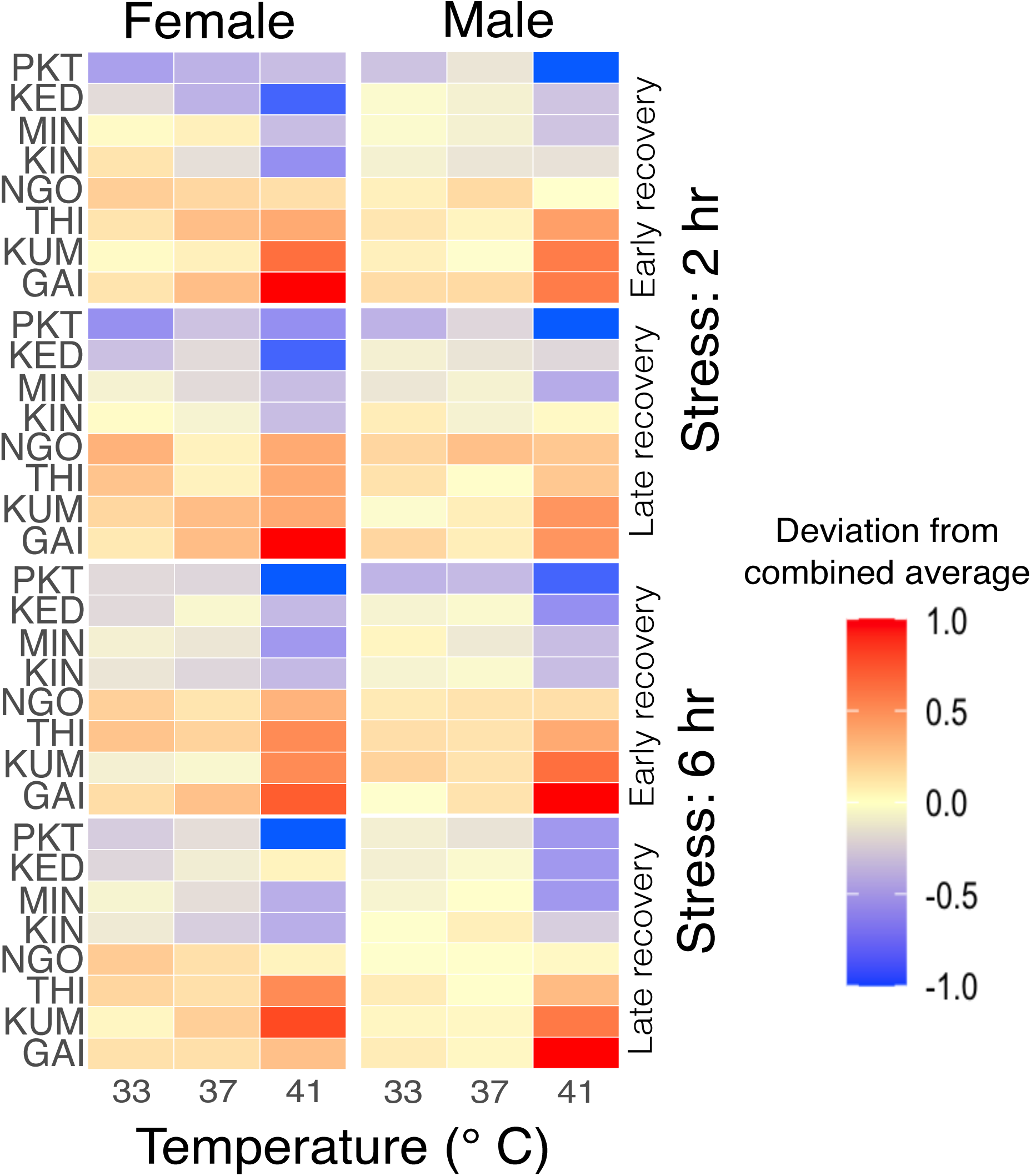
Deviation in adult *Aedes aegypti* survival in comparison to combined average following varied temperature exposures. Male and female adults were exposed to three extreme temperatures for 2 hours (Supplementary figures 1a and 1b) and 6 hours (Supplementary figures 1c and 1d). Following the exposure, the survival of the mosquitoes was evaluated after 2 hours (early) (Supplementary figures 1a and 1c) and 24 hours (later) (Supplementary figures 1b and 1d). For each treatment, 5 replicates were run, with 8 mosquitoes per replicate. The numbers within each heatmap tile represent the proportion survival of adult mosquitoes for a specified population, depending on the temperature treatment (Supplementary tables S2a and S2b). Figures were produced using R 4.2.3 and edited with Inkscape 1.3.

**Figure 2:**
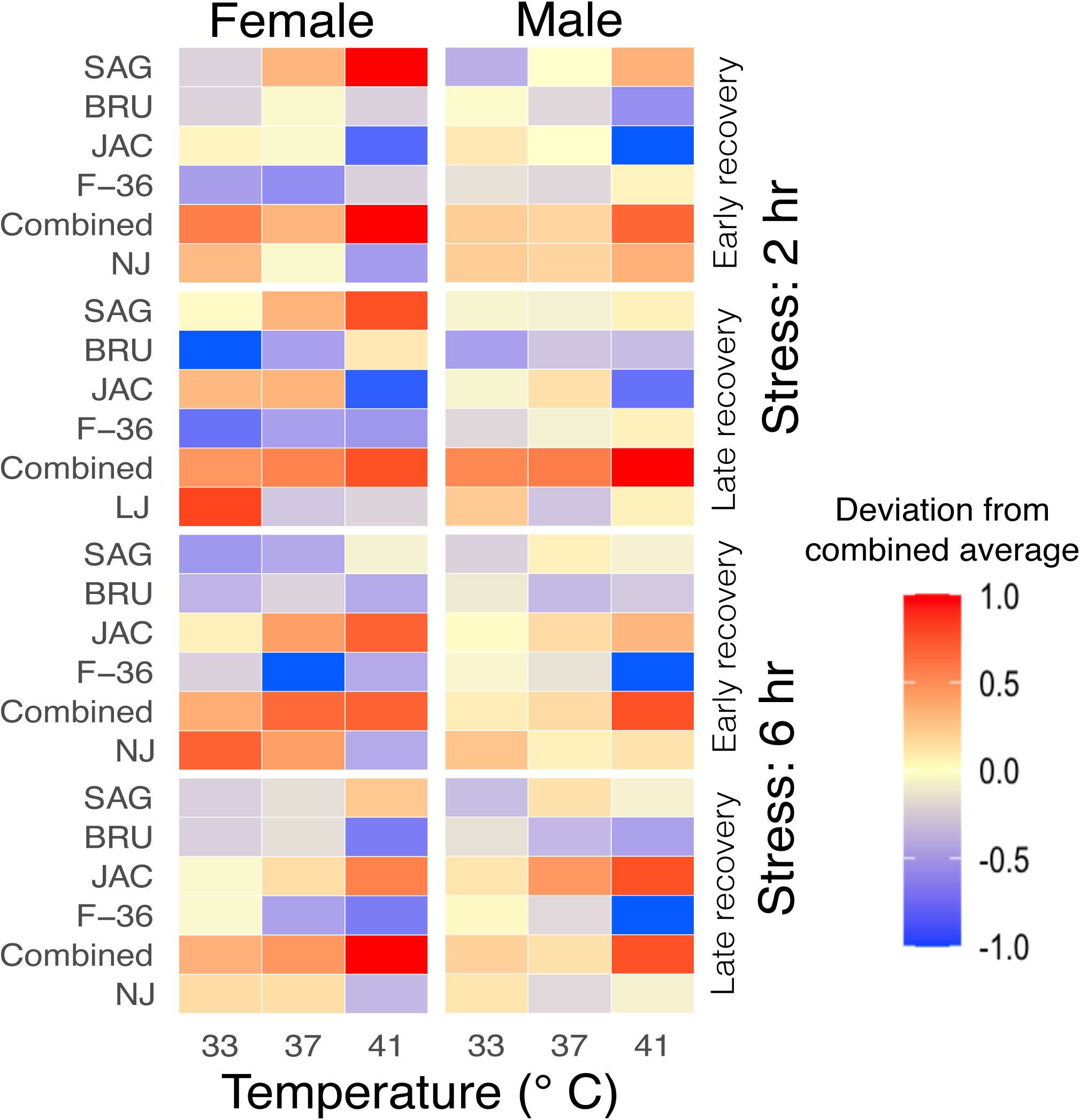
Deviation in adult *Aedes albopictus* survival in comparison to combined average following various levels of temperature exposure. Male and female adults *Ae. albopictus* following exposure to three distinct temperatures for 2 hours (Supplementary figures 1e and 1f) and 6 hours (Supplementary figures 1g and 1h). After these exposures, mosquito survival rates were assessed at 2-hour intervals (early) (Supplementary figures 1e and 1g) and 24-hour intervals (later) (Supplementary figures 1f and 1h). The experimental setup consisted of 5 replicates for each treatment, with each replicate comprising 8 mosquitoes. The numbers on each tile of the heatmap depict the proportion survival for that specific population of mosquitoes following the respective temperature treatment (Supplementary tables S2c and S2d). Figures were produced using R 4.2.3 and edited with Inkscape 1.3.

### Thermal tolerance of *Aedes* eggs

Substantial variation occurred in the egg hatching rate among the *Ae. aegypti* populations (F = 6.027, *p* = 6.62 × 10^-7^) (**Fig. 3**). As a general trend, a significant reduction in egg hatching was found as the temperature levels increased (F = 2146, *p* < 2.2 × 10^-16^) (**Supplementary figure 4a and 4b)**. Notably, the impact was more pronounced following the exposure to > 40 °C after the 2-hr stress period and above 37 °C after the 6-hr stress period (**Fig. 3, Supplementary table S4a**). Following exposure to 41 °C for 2-hr, the egg survival rates varied between 57.54±2.78% to 60.24±4.57% for the mosquito populations originating in areas where human population density is low (<55/km^2^), whereas the egg hatching varied between 66.53±3.38% to 78.07±2.76% for the mosquito populations collected from areas where human density is moderate (225-425/km^2^) to high (>2000/km^2^) (**Supplementary table S4b**). Significant differences were observed between populations collected from high (urban) and low human (rural) density areas when exposed to 45 °C for 6 hrs (F = 96.8, *p* < 2.2 × 10^-16^); the lowest egg hatching, 2.39±1.1% was encountered for a population, Kedougou (human density 12.25/km^2^), collected from rural habitats; and the maximum egg hatching was found for Ngoye population, 36.23±4.45%, a moderately human dense area (233/km^2^).

**Figure 3:**
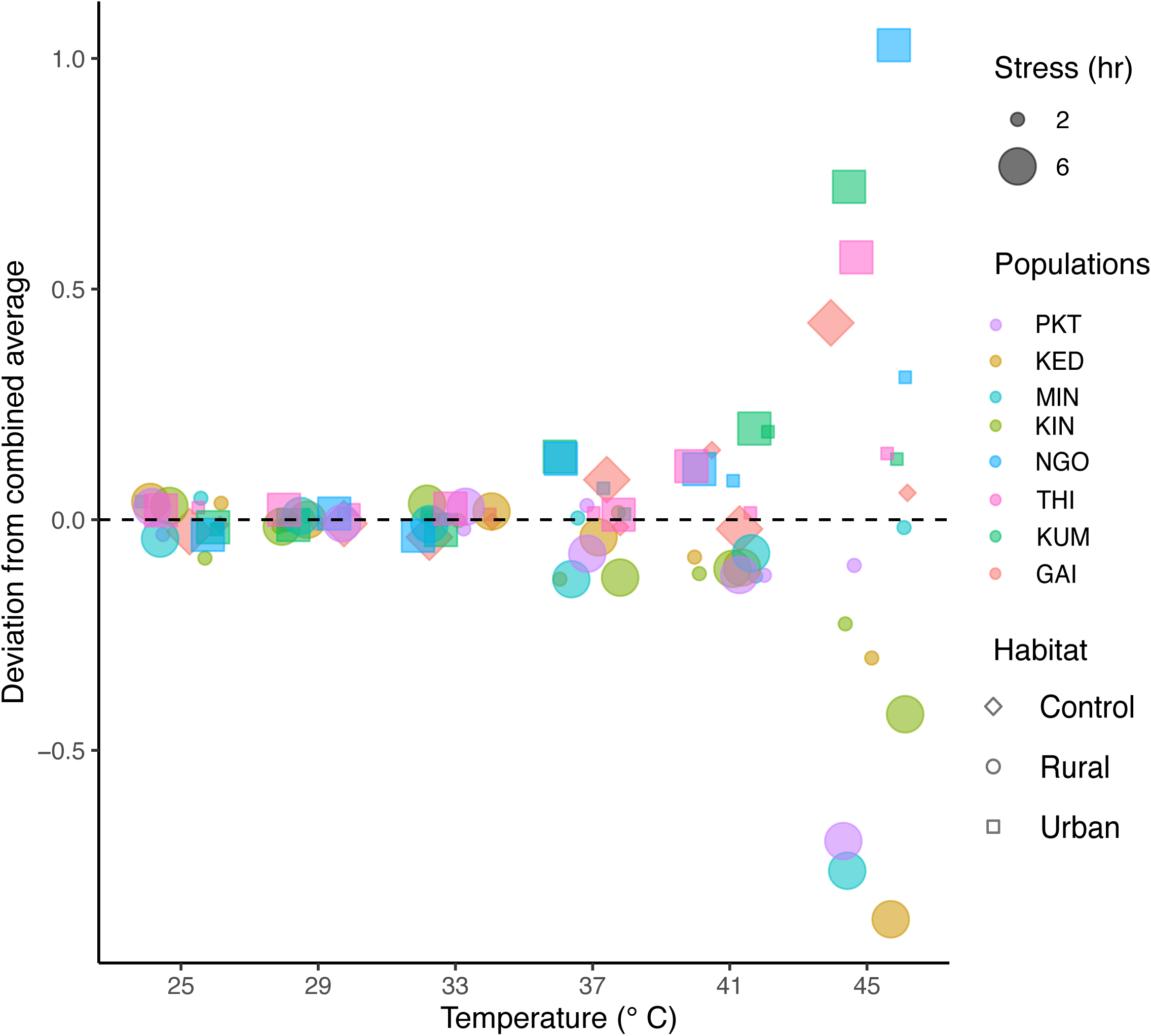
Deviation in egg hatching of *Aedes aegypti* populations in comparison to combined average following different temperature exposures. Egg hatching within the populations after exposure to experimental temperatures for 2 hours (Supplementary figure 4a) and 6 hours (Supplementary figure 4b). Each temperature-treatment combination was experimented with 10-12 replicates with 30-50 eggs. The numbers in each tile depict the proportion of hatching for the respective populations after each temperature exposure (Supplementary tables S4a and S4b). Figures were produced using R 4.2.3 and edited with Inkscape 1.3.

A significant decrease in egg hatching was observed among *Ae. albopictus* following elevated temperature stress (> 40 °C) compared to the *Ae. aegypti* (F = 40.59, *p* = 1.2 × 10^-16^) (**Supplementary figure 2c and 2d; Supplementary table S3b**). After exposure to 45°C, the egg-hatching rate for the lab-generated hybrid population “Combined” was 11.89±1.18% and 4.54±0.6% after 2-hr and 6-hr long temperature stress, respectively. Inter-populations differences in hatching rates occurred across populations under various temperature exposures (F = 0.665, *p* = 0.007) (**Fig. 4; Supplementary figure 4c and 4d; Supplementary tables S4c and S4d**). The lab-maintained population (NJ), the control group, exhibited higher hatching rates than other *Ae. albopictus* populations across all the temperature levels.

**Figure 4:**
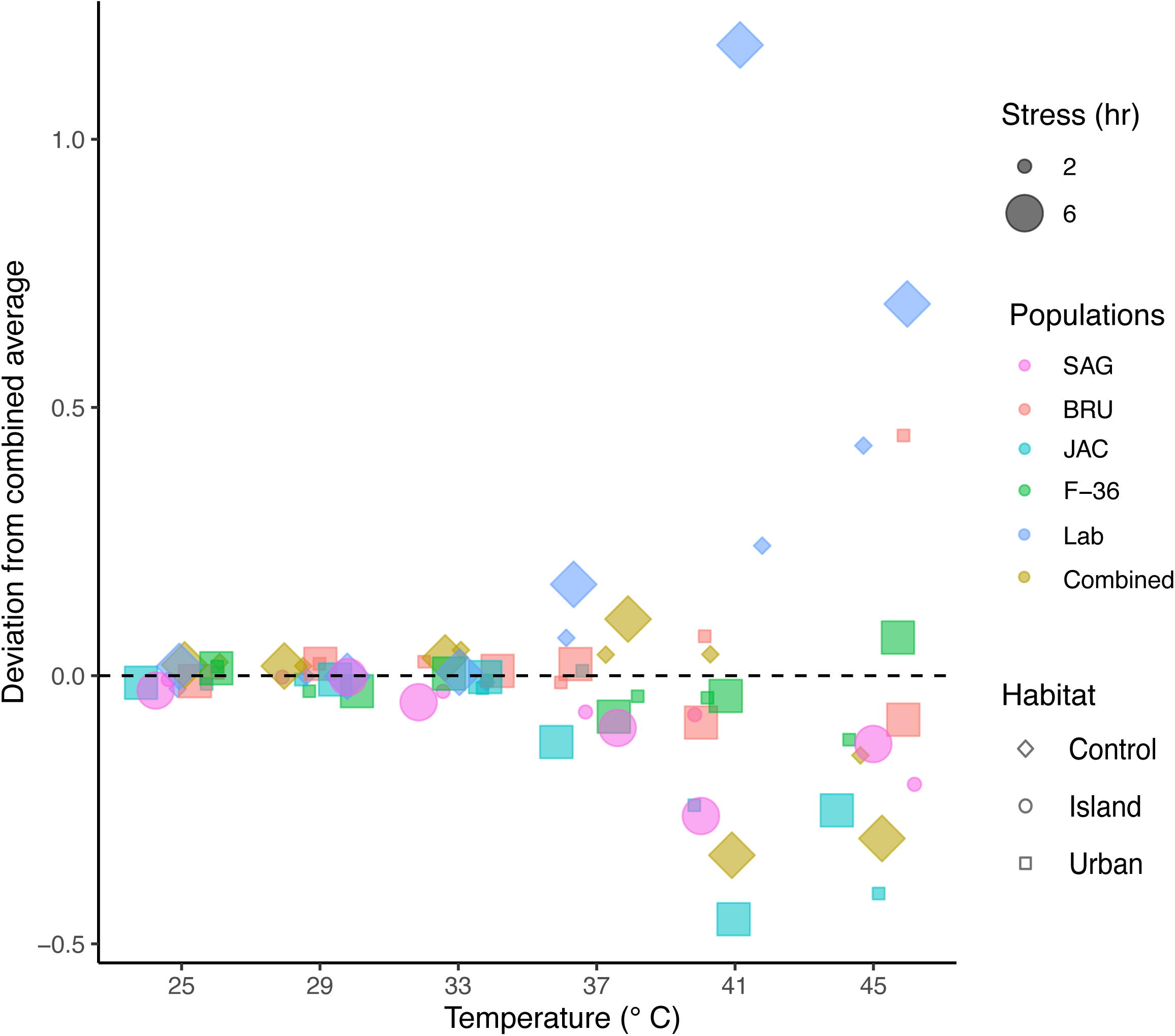
Deviation in egg hatching of *Aedes albopictus* populations in comparison to combined average following varying temperature exposures. Egg hatching in the *Aedes albopictus* populations occurred following exposure to varying temperature regimens for 2 hours (Supplementary figure 4c) and 6 hours (Supplementary figure 4d). Each experimental condition consisted of 10-12 replicates, each replicate having 30-50 eggs. The numerical values within individual tiles of the heatmap denote the proportion of egg hatching for the corresponding populations following each discrete temperature exposure (Supplementary tables S4c and S4d). Figures were produced using R 4.2.3 and edited with Inkscape 1.3.

### Influential factors impacting egg hatching success in *Ae. Aegypti*

In the initial phase of the tri-fold analysis, correlation analysis evaluated the relationships among ecological, behavioral, and ancestry indices. The strong correlation of 97% between the host preference and ancestral origin suggests an intertwined relationship between these two indices concerning egg thermotolerance (test statistic = 118.6215, degrees of freedom = 930, *p* < 2.2 × 10^-16^). Following the correlation test, a binomial logistic regression modeling approach evaluated the significant variables influencing egg hatching. We observed that, in addition to temperature and duration of stress, a comprehensive set of factors associated with specialization on human hosts and habitats (collectively: human specialization syndrome) significantly influenced the egg survival of the populations (**Fig. 5**).

**Figure 5:**
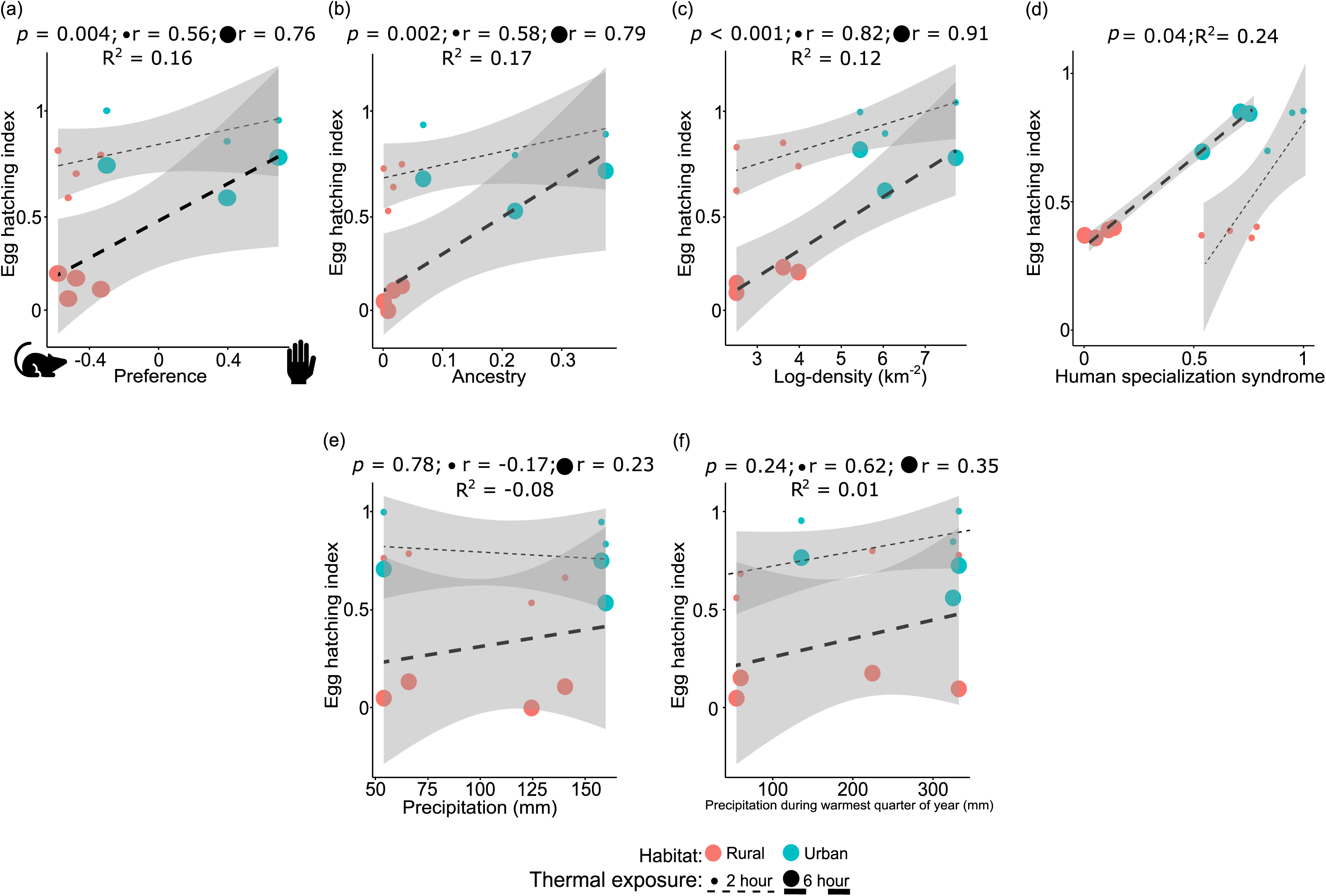
Factors influencing egg hatching success in *Aedes aegypti*. (a) The influence of host preference on egg hatching with a focus on the duration of thermal stress reveals a robust linear trend (*p* = 0.004). (b) The significant positive effect of ancestral origin on the egg-hatching success of *Ae. aegypti* mosquito eggs (*p* = 0.002). (c) The positive influence of density on egg hatching rate reveals that eggs originating from areas with higher human density exhibit higher hatching rates (*p* = 1.38 × 10^-8^). (d) Variables in (a), (b) and (c) can be combined in an index of human specialization syndrome, which accounts for 24% of variation in the final model ([LRT] *p* = 0.04). (e) Precipitation regimes (*p* = 0.78) and (f) precipitation during the warmest months of the year (*p* = 0.24) had no significant association with egg thermotolerance. ‘r’ values denote the correlation coefficient between the egg thermotolerance and the biological factors. R^2^ denotes the proportion of variance for egg thermotolerance associated with the specific biological factor(s). Figures were produced using R 4.2.3 and edited with Inkscape 1.3.

After evaluating the human specialization syndrome influencing egg hatching, we disentangled each factor as independently associated with egg thermotolerance. Temperature (Estimate = -0.31, *z* = -96.32, *p* < 2 × 10^-16^) and length of heat stress (Estimate = -0.19, *z* = -26.79, *p* < 2 × 10^-16^) showed a strong negative association with egg hatching, indicating higher temperatures for longer periods negatively impact egg hatching. The influence of host preference on egg hatching index demonstrated a strong linear fit (Estimate = 0.43, *z* = 2.91, *p* = 0.004, **Fig. 5a**). Within human specialist syndrome, the influence of ancestral origin emerged as a more prominent predictor for egg thermotolerance (Estimate = 1.6, *z* = 3.09, *p* = 0.002), accounting for ∼17% of the variability (**Fig. 5b**). Moreover, thermotolerance of egg survival was positively influenced by human density (Estimate = 0.0002, *z* = 5.68, *p* = 1.38 × 10^-8^), indicating that eggs originating from areas with higher human density have increased egg thermal tolerance (**Fig. 5c**). When assessing the significance of various factors that can explain egg thermotolerance, we observed that factors contributing to human specialist syndrome (host preference, human ancestry, and human population density) account for 24% of the egg thermotolerance differences (**Fig. 5d)**. Concurrently with temperature and stress conditions, when examining the impact of moisture availability parameters (precipitation [*p* = 0.78] and precipitation during the warmest months of the year [*p* = 0.24]), no significant association was found (**Fig. 5e and 5f**). Nevertheless, incorporating climatic factors alongside the human specialization syndrome results in a comparatively weak relationship (7% of variation in egg thermotolerance). Our results indicate a strong correlation between preference for human hosts and increased egg hatching rates under thermal stress in populations collected from densely populated urban habitats (**Supplementary figure 5**). Notably, the correlation between preference for human host and egg thermal tolerance was apparent when eggs were exposed to temperatures at or above 37°C. This underscores a clear and discernible impact of human specialization syndrome on egg hatching during thermal exposure of mosquitoes collected from two urban and rural habitats. As the temperature level and duration of thermal exposure increased, a more pronounced influence of human preference on the egg hatching index became evident (**Supplementary figure 5**). Overall, our findings show a robust association between egg hatching of *Ae. aegypti* and temperature stress, human specialist ancestry, and human density. However, there was not a definite relationship between the thermotolerance of adult mosquitoes and ecological, behavioral, and ancestry indices. Sex-specific studies revealed only adult female survival was moderately influenced by host preference, but few of the conditions are signfiicant (**Supplementary figure 6**). However, this was insignificant and would require more populations known host preference to determine a potential relationship.

## Discussion

Understanding thermal performance concerning the life history traits of mosquitoes has provided invaluable insight into their thermal adaptations (Couper et al., 2021; Holmes & Benoit, 2019; Lahondère & Bonizzoni, 2022; Mordecai et al., 2019). The increase in anthropogenic activities is likely to have implications for the biology of specific mosquitoes with their response to thermal stress (Rochlin et al., 2016; Schrama et al., 2020). The thermal tolerance of eggs has not been extensively examined. We identified that urban-associated mosquito populations exhibited a notably higher egg hatching rate than rural-associated mosquito populations following thermal stress. Furthermore, our analysis of ecological and behavioral indices indicated that egg thermal tolerance is a previously unrecognized part of human specialization syndrome. This thermal tolerance is robustly influenced by the key factors associated with the syndrome (host preference, genomic ancestry, and human density). Interestingly, our findings suggest that climatic variables do not exert a discernible impact on the thermal tolerance of *Ae. aegypti* eggs. In contrast, adult thermal tolerance showed no significant differences or discernible trends among the populations for both *Ae. aegypti* and *Ae. albopictus*. Our study provides a foundation to determine the intricate interplay between ecological factors, heat stress, and mosquito egg thermal tolerance.

Assessing adult mosquito survival under different temperatures provides insight into their thermal tolerance and adaptability (Ware-Gilmore et al., 2023; Zani et al., 2005). In this study, several populations of *Ae. aegypti* and *Ae. albopictus* were characterized to evaluate their temperature tolerance and assess variations in survival. Overall, we found variations in survival rates between males and females, with a consistent decline in survival with temperature increases. However, we detected only a few significant differences in survival between mosquito populations collected from rural and urban habitats, particularly among the female *Ae. aegypti*. We did note a trend where populations from urban areas have a generally higher thermal tolerance than those from rural areas, but this was not significant in relation to factors such as ancestry or human population density. It is likely that if *Ae. aegypti* was collected over a larger geographic range or substantially more populations were collected more differences would have been observable in adults, as seen in *Ae. albopictus* (Carlassara et al., 2024). Interestingly, no significant differences in survival were observed between tropical and temperate *Drosophila melanogaster*, residing within the experimental thermal range of 11 to 32 °C (Trotta et al., 2006). The fruit flies exhibited thermal plasticity in developmental rate, body size, and fertility; however, distinct adaptive responses following heat stress were not prominent among flies from different origins (Trotta et al., 2006). The survival of an organism in nature is contingent upon factors such as season and geographical location (Mitrovski & Hoffmann, 2001; Rosewell & Shorrocks, 2008), and laboratory measurements may not align with survival in the wild (Mołoń et al., 2020), as adults can effectively behaviorally thermoregulate depending upon the environmental conditions to moving to a more favorable microhabitat. Our study also found that *Ae. albopictus* has lower thermal tolerance than *Ae. aegypti*. Notably, *Ae. albopictus* displays a remarkable ability to endure colder climates compared to *Ae. aegypti* (Reinhold et al., 2018). Our findings are consistent with a previous study on *Aedes*, which demonstrated a prevalence of *Ae. aegypti* in urban locales contrasted with *Ae. albopictus* is more common in rural regions of Thailand (Tsuda et al., 2006). This distribution difference suggests that, *Ae. aegypti* has an enhanced ability to tolerate heat when compared to *Ae. albopictus* could be a factor that allows this mosquito to persist in urban areas in tropical and dry regions. For both of the studied species, we consistently demonstrated that under various levels of heat stress, female mosquitoes exhibited higher survival rates compared to male mosquitoes, a pattern commonly observed in arthropods (Bodlah et al., 2023; Chen et al., 2018; Weaving et al., 2022). In addition to physiological and behavioral distinctions, thermoregulatory behavior, such as engaging in evaporative cooling during the blood-feeding process to counter the rapid temperature increase, may contribute to the sex-specific difference in survival (Benoit et al., 2011; Kingsolver & Huey, 2008; Lahondère & Lazzari, 2012; Nylin & Gotthard, 1998; Reinhold et al., 2021). Furthermore, the observed phenomenon may also be influenced by the size disparity between female and male mosquitoes. Considering that female mosquitoes are slightly larger than their male counterparts, these size differences are also likely to play an important role in shaping survival patterns.

The egg stage cannot move to more favorable conditions, and exposure to high temperatures during the egg stage has been shown to increase mortality and reduce growth in subsequent development stages in insects (Potter et al., 2011). Among other insects, no differences in thermal tolerance were noted in the eggs of the tobacco hornworm, *Manduca sexta* (Potter & Woods, 2012) and there has been very little focus on this topic in other arthropod systems (Ajayi et al., 2023; Denlinger & Yocum, 2019; Harvey et al., 2020; Hilker et al., 2023). Our study revealed significant differences in the egg-hatching rates for *Ae. aegypti* based on their origin of collection (areas with high and low human population density) when exposed to temperature stress. Microclimatic variations within urban and rural environments, dry season intensity, and precipitation regimes can impact mosquito egg survival following heat stress (Mogi et al., 1996). The contrasting differences in the survival of mosquito eggs from urban and rural settings suggest mosquitoes that reside in more urbanized settings deposit eggs with increased heat tolerance (Lim et al., 2021; Simon et al., 2020). A sparsely vegetated urban site can reach temperatures up to 11°C higher than the surrounding countryside (Chen et al., 2006; Mohajerani et al., 2017), due to increased concrete and asphalt surfaces along with other human activities (Oke, 1982; Shashua-Bar & Hoffman, 2000). Also, the variability in available materials for mosquito egg deposition across different habitats significantly contributes to the observed differences in egg hatching among collection sites. Thermally conductive materials such as rubber, plastic, concrete and earthenware, are abundant in a densely populated area, and mosquitoes prefer these substances for egg laying (Seng & Jute, 1994). In contrast, in wild areas or rural areas with fewer humans mosquitoes typically opt for tree holes or natural habitats to lay eggs (Suganthi et al., 2014). Notably, during hot summer days or exposure to direct sunlight, the live wood remains cooler due to multiple processes (e.g. evaporative cooling) compared to other substances (Ellison et al., 2017; Moss et al., 2019; Souch & Souch, 1993). Moreover, driven by the increased opportunity of biting human hosts and the availability of alternative water sources, mosquitoes tend to move towards densely populated habitats if there is a preference for humans (Chu et al., 2022; Fernández-Grandon et al., 2015; Valtonen et al., 2020). This shift necessitates an increased environmental stress tolerance, as seen in other terrestrial arthropods shifting to urban areas (Diamond et al., 2018; Harvey et al., 2020, 2023). Overall, we show that heat stress, specifically above 37 °C, significantly reduced *Aedes* egg hatching, and the population from urban areas with increased human preference demonstrated greater thermal tolerance compared to counterparts from more rural areas with a reduced human preference.

Alongside temperature stress, the interplay of urbanization, microclimate variation, and other potential ecological factors are likely to have an influential role in shaping the survival of the populations (Erraguntla et al., 2021; Flores Ruiz et al., 2022; Murdock et al., 2017; Rochlin et al., 2016; Wang et al., 2020). Genetic diversity and habitat adaptation greatly influence the biology of mosquitoes (Couper et al., 2021; McBride et al., 2014). In the African landscape, mosquito populations display admixture between the domestic human-specialist form (*‘Aaa’*) and forest form (*‘Aaf’*), which is the prevailing pattern of genetic structure among the *Ae. aegypti* populations (Rose et al., 2020). The proportion of *‘Aaa’*-like ancestry (admixture) is the main driver that influences traits such as host preference and underscores the complex evolutionary forces that play roles in human interaction and the successful exploitation of ecological niches by mosquitoes (Rose et al., 2023). Although *‘Aaa’* is traditionally associated with domestic habitats, not all *‘Aaa’* populations are necessarily urban, and not all urban populations are heavily skewed towards ‘*Aaa’* form. Populations from the highly human-dense urban area of Kumasi (Ghana), have a genetic background that is predominantly *‘Aaf’*, but with substantially more ‘*Aaa’* ancestry than is observed in nearby rural populations (Rose et al. 2020). Interestingly, our study indicates a high thermal tolerance in the eggs from the Kumasi population. Given the overall association of *‘Aaa’*- like ancestry with egg thermal tolerance, this could reflect increased ‘*Aaa’* ancestry in specific loci controlling thermal tolerance relative to genome-wide levels, or could reflect variation in thermal tolerance within ‘*Aaf’*, beyond the differences observed between the two forms. Our study supports the previous research that suggests genetic variations and local adaptations influence the physiological responses of insects to temperature fluctuations (González-Tokman et al., 2020; Hoffmann & Sgrò, 2011; Sternberg & Thomas, 2014). The link between ancestry and thermal tolerance suggests that higher thermal tolerance of eggs is likely a physiological trait, along with an increased preference for humans, that has allowed an anthropophilic shift from a more generalist nature.

Temperature resistance in *Ae. albopictus* eggs are notably diminished compared to *Ae. aegypti* eggs corroborate previous findings (Juliano et al., 2002). All the *Ae. albopictus* populations in this study originated from regions of high human density and similar geo-climatic conditions, with only one exception, which originated from a varied habitat on Kyushu Island, Japan (Mushegian et al., 2021). Surprisingly, despite the diverse origin of the Kyushu Island populations, the egg-hatching response closely aligns with the other populations originating from the East Coast, USA. This similarity may be attributed to a combination of factors, including the increased frequency of hotter days on the island due to climate change, coupled with amplified atmospheric moisture, which results in intense precipitation events (Kawase et al., 2019; Takabatake & Inatsu, 2022) leading to increased mosquito epidemics (Mogi, 1996; Yang et al., 2021). Kyushu Island is home to three container terminals that primarily import cargo, and it has been reported to have a much higher mosquito prevalence (Yang et al., 2021). Within the terminal areas, the mosquito oviposits in enclosed habitats and the swift progression of climate change on the island may be fostering the thermotolerance of the eggs. Prolonged laboratory adaptation might contribute to the blending effect of differential thermotolerance in eggs (Hoffmann & Ross, 2018; Ross et al., 2019), as most of the *Ae. albopictus* lineages have been in laboratory settings for at least 15 generations. Recent population genetics study of *Ae. albopictus* from several parts of the globe confirmed that the USA lineage is derived from Japan (Garcia-Rejon et al., 2021). Several migration events (local and long-distance dispersal, regional gene flow) influence the invasion and successful colonization of *Ae. albopictus* mosquitoes in the East Coast of the USA (Gloria-Soria et al., 2022; Vega-Rúa et al., 2020). Further investigations are warranted to elucidate the specific mechanisms underlying this unexpected variation in temperature tolerance among geographically distinct populations.

Our study represents one of the first extensive analyses of temperature stress on *Aedes* mosquito adults and eggs of distinct origin and how genetic variations, local adaptations, or other factors influence the physiological responses of *Ae. aegypti* eggs to temperature fluctuations. We found that the egg-hatching rates were positively impacted by increasing human density, which suggests that egg thermotolerance could be critical for mosquito survival during climate change and rural-to-urban transitions. The process of urbanization, encompassing rapid urban growth, unplanned expansion, and human population density, unquestionably shapes mosquito populations (Carlassara et al., 2024; Kolimenakis et al., 2021), leading to differential resilience of mosquito populations in the face of abiotic extremities. Anthropogenic changes frequently lead to the transformation of natural landscapes, creating more breeding sites for mosquitoes (Wilke et al., 2019). The artificial containers and inadequately maintained water storage in urban environments favor mosquito breeding, even during dry periods (Flaibani et al., 2020; Getachew et al., 2015; Rose et al., 2020). These artificial water containers are commonly warmer than those found in a rural setting, highlighting that deposited eggs require increased thermotolerance, as demonstrated in this study. In contrast, mosquitoes oviposit in tree holes are buffered from the effect of high temperature, as wood is a living tissue, high in water content, that will not overheat in the sun or on a hot day. Our findings contribute to the growing evidence that insect life stages, particularly eggs, exhibit adaptive tolerance to global increases in temperature up to a specific limit (Carlassara et al., 2024; Couper et al., 2023; Quan et al., 2023; Reinhold et al., 2018). Further research is warranted to elucidate the specific physiological and molecular mechanisms underlying the observed differences in egg survival, which will contribute to our understanding of the adaptive potential of mosquito populations with climate change.

## FIGURE LEGENDS

**Supplementary Figure 1: Adult survival of *Aedes aegypti* and *Aedes albopictus* populations following thermal stress**

Male and female adults of *Ae. aegypti* (a-d) and *Ae. albopictus* (e-h) were exposed to varying levels of temperatures for 2 hours (a, b, e, and f) and 6 hours (c, d, g, and h). Following the thermal exposure, adult survival was evaluated after 2 hours (a, c, e, and g) and 24 hours (b, d, f, and h). Tests were carried out on eight (8) *Ae. aegypti* populations and six (6) *Ae. albopictus* populations. Figures were produced using R 4.2.3 and edited with Inkscape 1.3.

**Supplementary Figure 2: Comparative responses of adult survival and egg hatching of *Aedes aegypti* and *Aedes albopictus* following temperature-induced stress**

Laboratory-maintained populations of *Aedes aegypti* (Gainesville: GAI) and *Aedes albopictus* (New Jersey: NJ) were experimented with adult mosquito survival (a and b) and egg hatching (c and d) after temperature stress. A 2-hour (a) and 6-hour (b) stress on adult survival and 2-hour (c) and 6-hour (d) stress on egg survival between the two populations were conducted. *p* values represent a significant difference among the populations following respective treatment. Figures were produced using R 4.2.3 and edited with Inkscape 1.3.

**Supplementary Figure 3: Comparison of adult male and female survival under thermal stress in *Aedes aegypti* and *Aedes albopictus***

The contrast in adult survival between laboratory-maintained populations of *Aedes aegypti* (Gainesville: GAI) and *Aedes albopictus* (New Jersey: NJ) male and female mosquitoes following thermal stress across three different temperature levels. The bar plot depicts the mean proportion of surviving mosquitoes, with error bars representing standard errors. Figures were produced using R 4.2.3 and edited with Inkscape 1.3.

**Supplementary Figure 4: Egg hatching of *Aedes aegypti* and *Aedes albopictus* populations following temperature exposures**

Egg hatching within eight *Ae. aegypti* populations (a and b) and six *Ae. albopictus* populations (c and d) after exposure to varying levels of thermal stress for 2 hours (a and c) and 6 hours (b and d). Figures were produced using R 4.2.3 and edited with Inkscape 1.3.

**Supplementary Figure 5: Correlation between human preference and egg-hatching**

Elevated temperatures and extended periods of thermal stress contribute to more pronounced changes in the association between human preference and egg-hatching. ‘r’ values represent correlation coefficient between egg hatching and host preference, R^2^ denotes the proportion of variance for egg hatching that is explained by the host preference. *p* values indicate a significant influence of host preference on the egg hatching of the mosquito populations. Figures were produced using R 4.2.3 and edited with Inkscape 1.3.

**Supplementary Figure 6: Relationship of host preference and *Aedes aegypti* adult thermotolerance**

(a) No influence was found between host preference and adult thermotolerance. Sex-specific analysis revealed (b) only female survival moderately influenced by host preference, (c) although male survival shows no discernible impact from host preference. *p* values and * denotes a significant influence of host preference on the adult thermotolerance. ‘r’ values represent correlation coefficient between the adult thermotolerance and host preference and R^2^ denotes the proportion of variance for adult thermotolerance that is explained by host preference. Figures were produced using R 4.2.3 and edited with Inkscape 1.3.

**Supplementary Table 1: Information on mosquito populations**

Collection localities and information about the *Aedes aegypti* and *Ae. albopictus* populations. Human population densities are calculated within a 20km radius. Latitude and Longitude are the coordinates of egg ovitraps from collection areas.

**Supplementary Table 2: Adult exposure to temperature stress**

Statistical significance of temperature treatment combinations on adult *Aedes aegypti* (**S2a**) and *Ae. albopictus* (**S2c**) survival. The average survival of *Ae. aegypti* (**S2b**) and *Ae. albopictus* (**S2d**) adults following several temperature-treatment combinations.

**Supplementary Table 3: Relative comparison of adult survival and egg hatching of lab-maintained *Aedes aegypti* and *Ae. albopictus* populations. Related to Supplementary figure 2.**

Average survival of laboratory-maintained populations of *Aedes aegypti* (Gainesville: GAI) and *Aedes albopictus* (New Jersey: NJ) adults (**S3a**) and eggs (**S3b**) following several temperature stresses.

**Supplementary Table 4: Egg exposure to temperature stress**

Statistical significance of temperature treatment combinations on *Aedes aegypti* (**S4a**) and *Ae. albopictus* (**S4c**) eggs hatching. The average egg hatching of *Ae. aegypti* (**S4b**) and *Ae. albopictus* (**S4d**) following temperature treatments.

## Acknowledgements

This study was partially supported by the National Institute of Allergy and Infectious Diseases of the National Institutes of Health under Award Number R01AI148551 and R21AI166633 (to J.B.B.). The following reagent was obtained through BEI Resources, NIAID, NIH: *Aedes albopictus*, Strain Gainesville, MRA-804, contributed by Sandra A. Allan. The following reagent was provided by Centers for Disease Control and Prevention for distribution by BEI Resources, NIAID, NIH: *Aedes albopictus*, Strain ATM-NJ95, Eggs, NR-48979.

## CRediT authorship contribution statement

**Souvik Chakraborty:** Conceptualization, Methodology, Software, Validation, Formal analysis, Investigation, Resources, Data curation, Writing – original draft, Writing – review & editing, Visualization, Supervision, Project administration.

**Emily Zigmond:** Formal analysis, Investigation, Resources, Data curation,Writing – review & editing.

**Sher Shah:** Formal analysis, Investigation, Resources, Data curation, Writing – review & editing.

**Massamba Sylla:** Resources.

**Jewelna Akorli:** Resources.

**Sampson Otoo:** Resources.

**Noah H. Rose:** Writing – Resources, review & editing.

**Carolyn S. McBride:** Writing – review & editing.

**Peter A. Armbruster:** Writing – Resources, review & editing.

**Joshua B. Benoit:** Conceptualization, Methodology, Validation, Investigation, Resources, Data curation, Writing – review & editing, Supervision, Project administration, Funding acquisition. **Declaration of competing interest**

The authors declare no conflicts of interest.

