## Supplemental Figures for "Thermal tolerance of mosquito eggs is associated with urban adaptation and human interactions"

Supplementary Figure 1: Adult survival of *Aedes aegypti* and *Aedes albopictus* populations following thermal stress

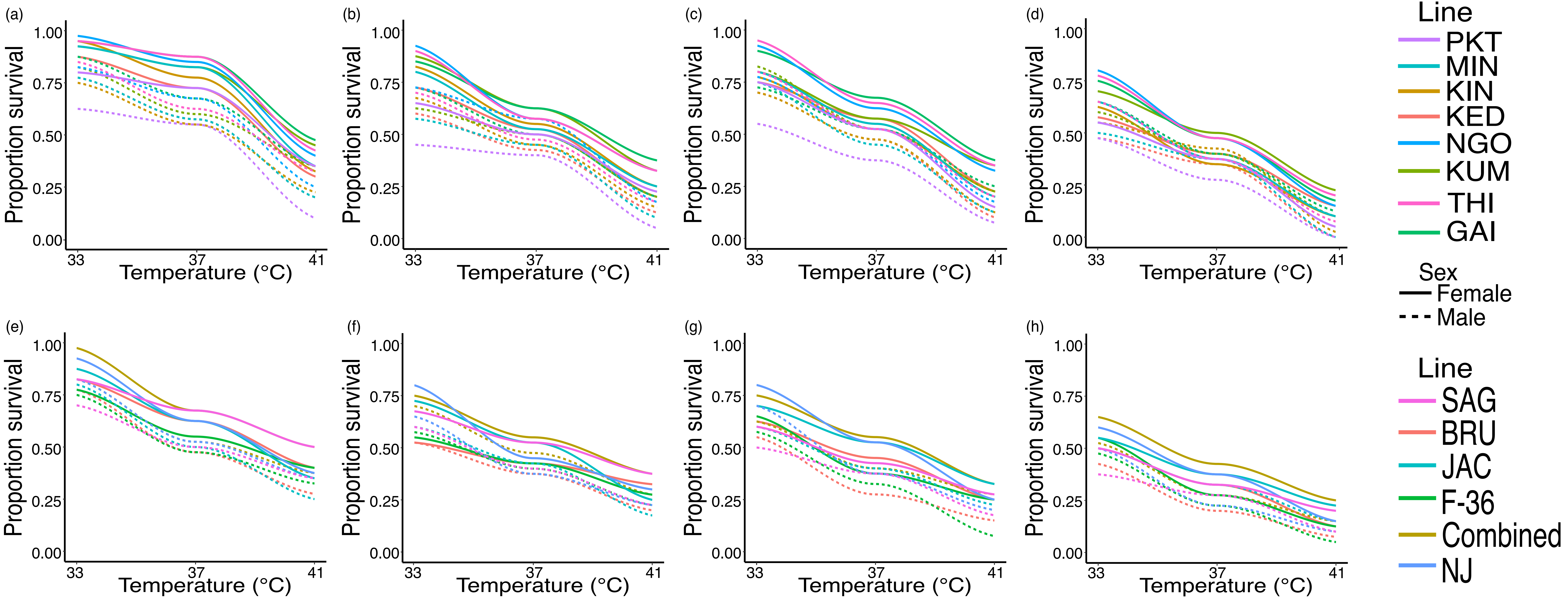

Supplementary Figure 2: Comparative responses of adult survival and egg hatching of *Aedes aegypti* and *Aedes albopictus* following temperature-induced stress

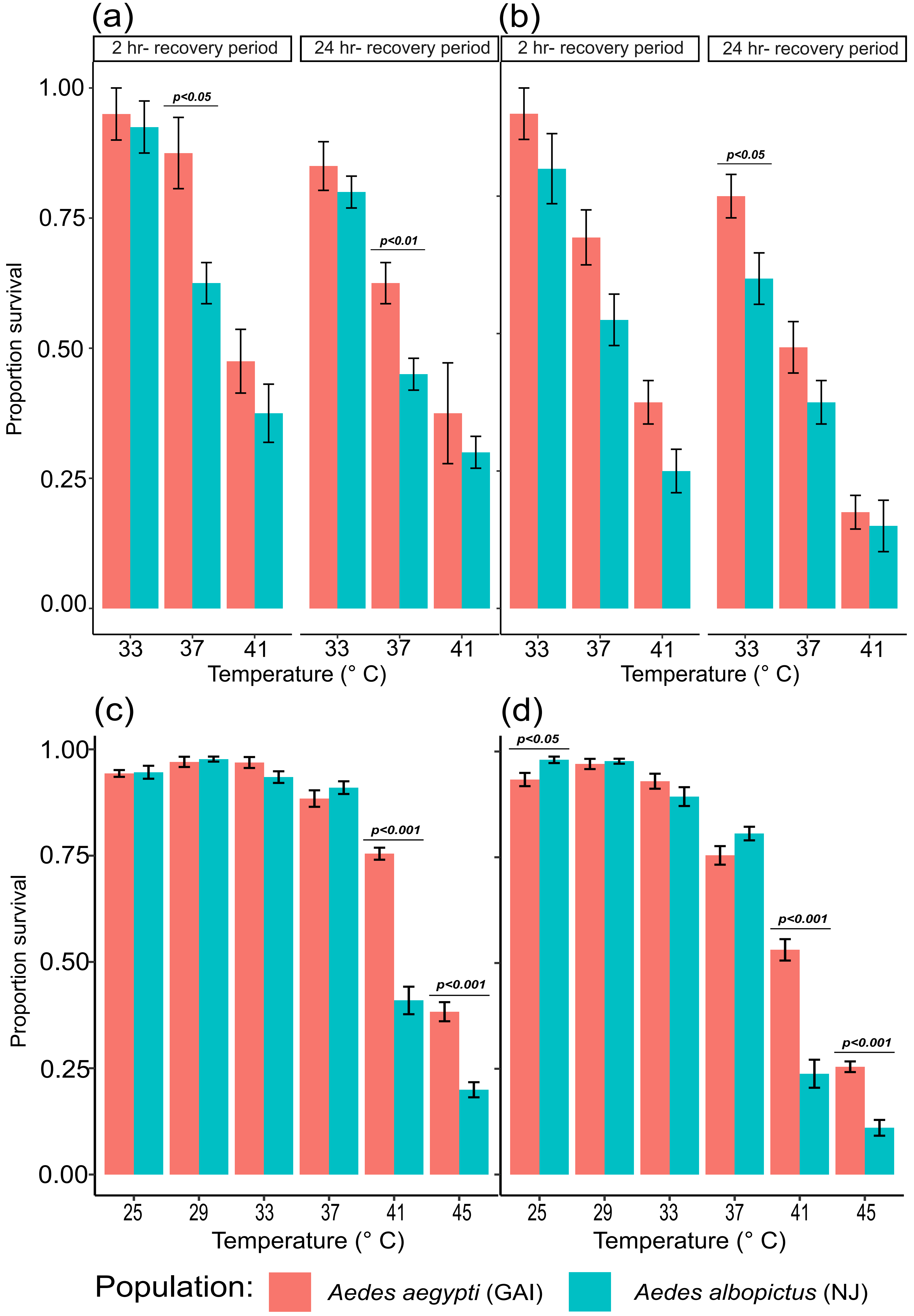

Supplementary Figure 3: Comparison of adult male and female survival under thermal stress in *Aedes aegypti* and *Aedes albopictus*

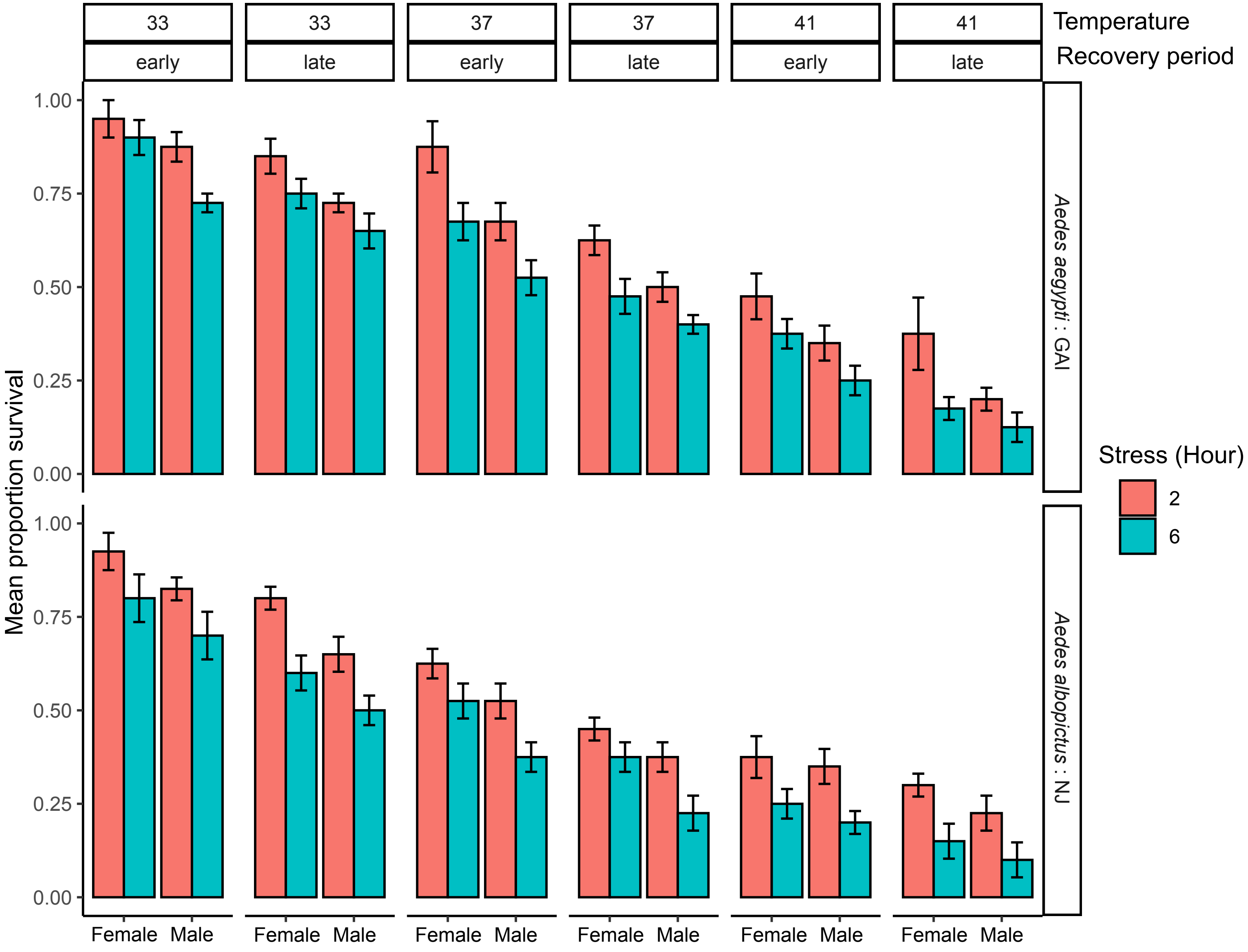

Supplementary Figure 4: Egg hatching of *Aedes aegypti* and *Aedes albopictus* populations following temperature exposures

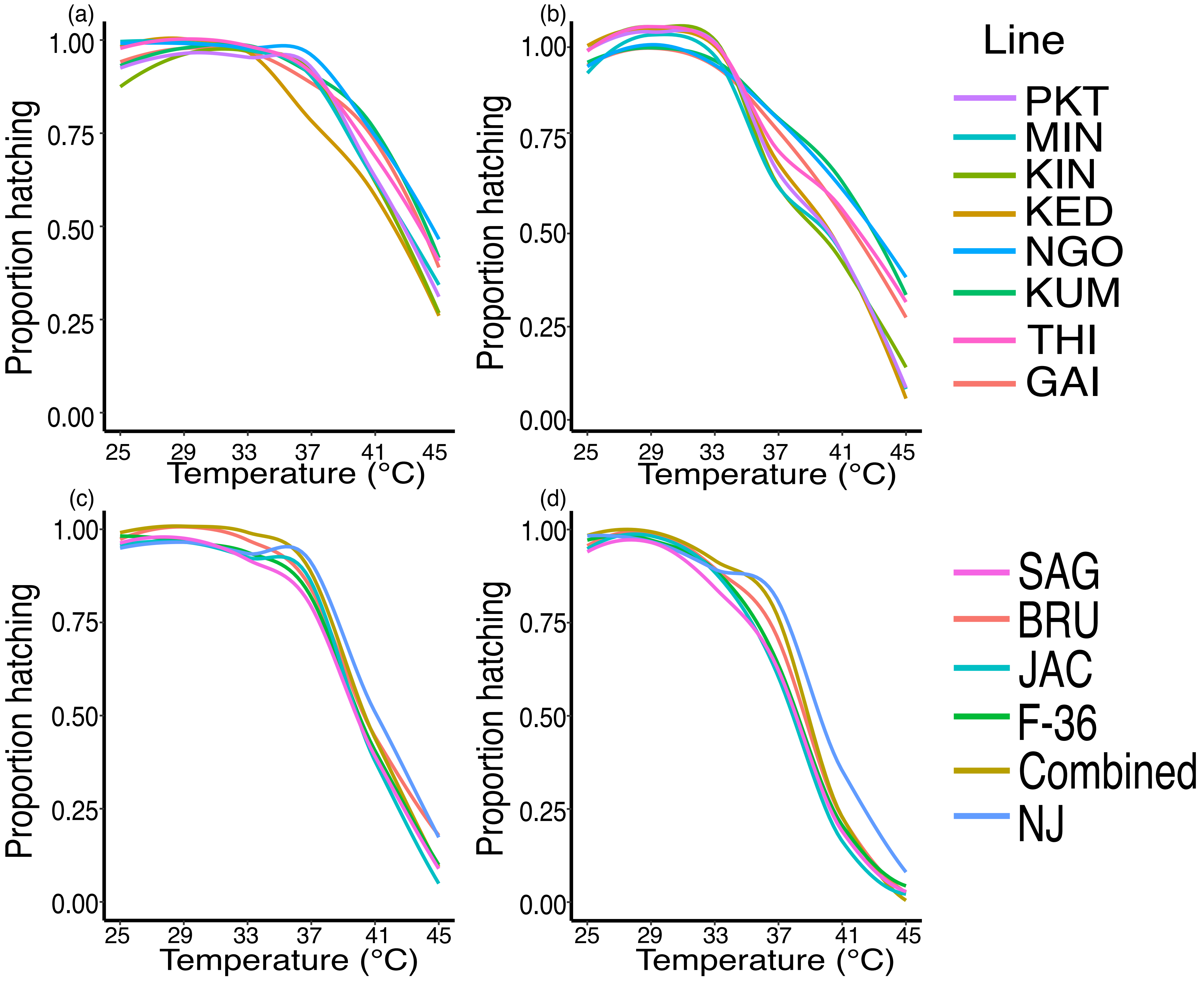

Supplementary Figure 5: Correlation between human preference and egg-hatching

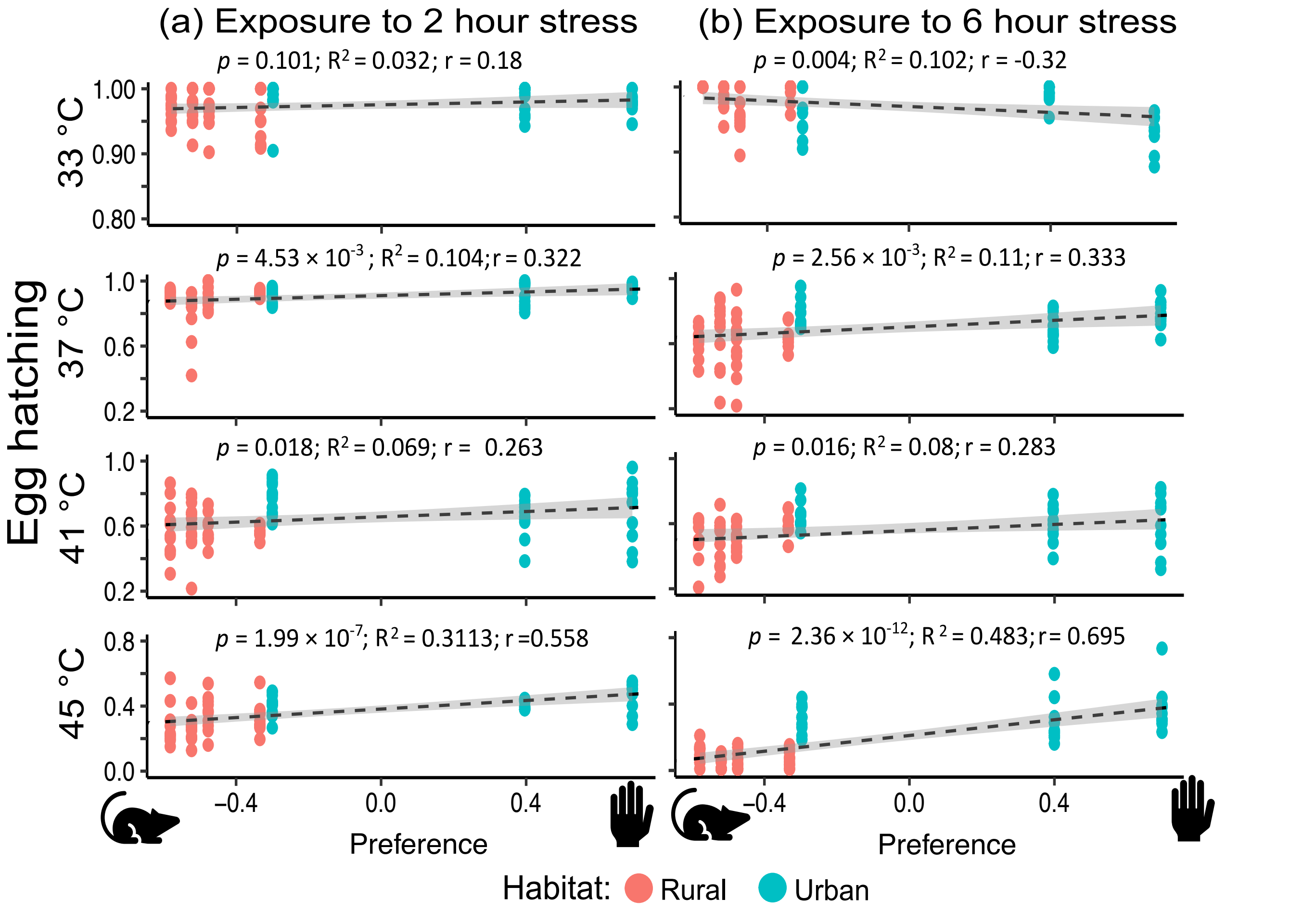

Supplementary Figure 6: Relationship of host preference and *Aedes aegypti* adult thermotolerance

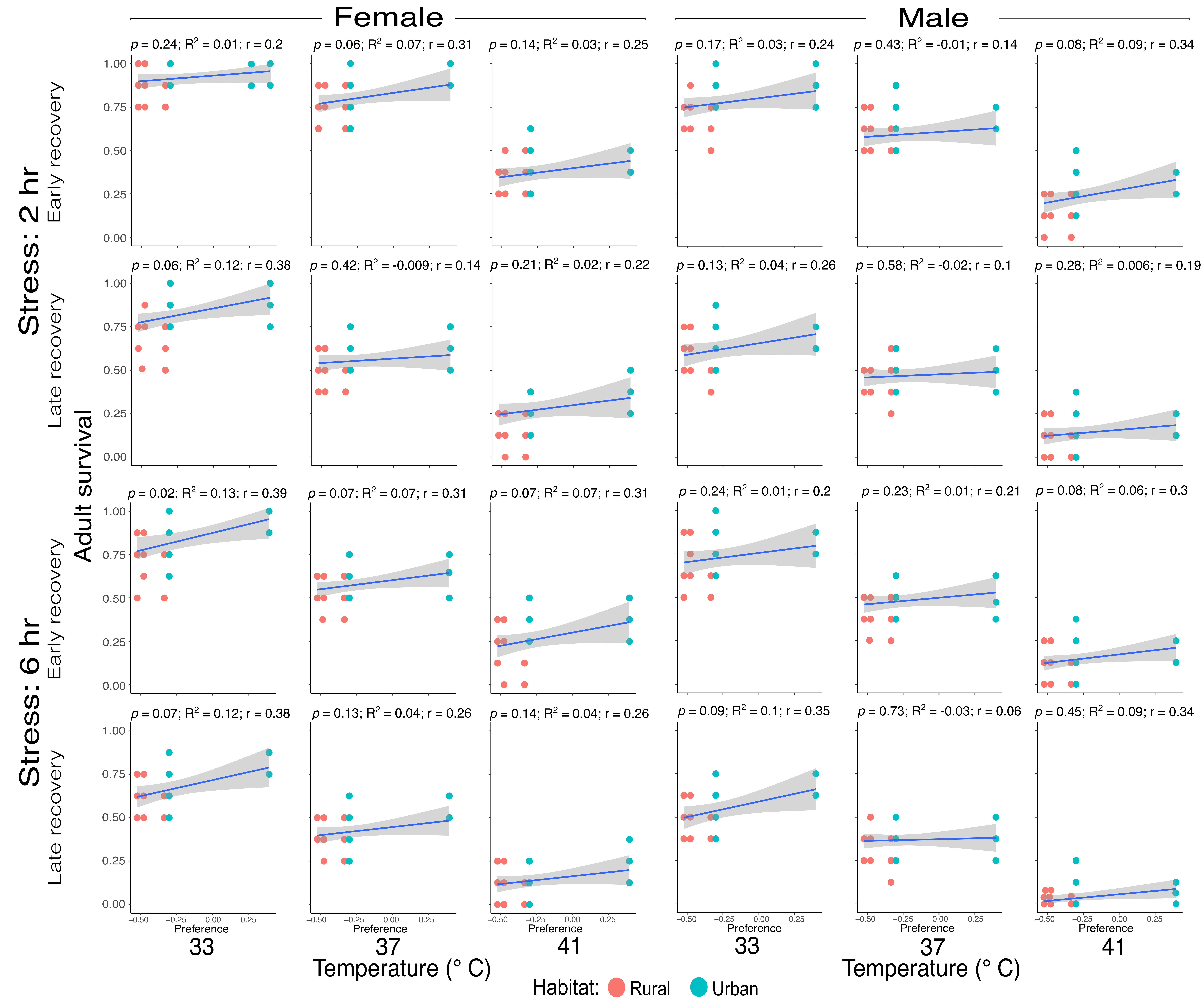
